# Histone lysine demethylase inhibition is a disease-modifying therapy for hypertrophic cardiomyopathy

**DOI:** 10.64898/2026.08.07.743611

**Authors:** Manjeet Singh, Yan Fan, Damir Alzhanov, Lingling Duan, Tram Anh Tran, Dinesh Ravindra Raju, Jinhua Wen, Christian Lopez Escobar, Matthias Peltz, Pietro Bajona, Xing Chao, Jun Liao, Dian J. Cao, Eric N Olson, Elisabeth D Martinez, Zhi-Ping Liu

**Affiliations:** Department of Internal Medicine-Cardiology, UT Southwestern Medical Center, Dallas, TX 75390, USA; Department of Bioengineering, The University of Texas at Arlington, Arlington, TX 76019, USA; Department of Molecular Biology, UT Southwestern Medical Center, Dallas, TX 75390, USA; Department of Pharmacology, UT Southwestern Medical Center, Dallas, TX 75390, USA; Eugene McDermott Center for Human Growth and Development, UT Southwestern Medical Center, Dallas, TX 75390, USA; Department of Internal Medicine-Histology core, UT Southwestern Medical Center, Dallas, TX 75390, USA; Lyda Hill Department of Bioinformatics, UT Southwestern Medical Center, Dallas, TX 75390, USA; Peter O’Donnell School of Public Health, UT Southwestern Medical Center, Dallas, TX 75390, USA

## Abstract

**Rationale:** Hypertrophic cardiomyopathy (HCM) is a common inherited cardiac disorder characterized by cardiac hypertrophy, fibrosis, arrhythmias, and sudden cardiac death (SCD). Although current therapies primarily target sarcomere dysfunction, the contribution of epigenetic dysregulation to HCM pathogenesis and its therapeutic potential remain poorly understood.

**Objective:** To determine whether pharmacological inhibition of histone lysine demethylases (KDMs) with JIB-04 can prevent or reverse HCM progression and to identify the underlying epigenetic mechanisms.

**Methods and Results:** We evaluated the pan-KDM inhibitor JIB-04 in Myh6^R403Q/+^ mice carrying the murine equivalent of the pathogenic human MYH7 R403Q mutation. JIB-04 prevented disease progression, reduced cardiac hypertrophy and fibrosis, preserved cardiac function, and completely prevented SCD in cyclosporin A– accelerated HCM. JIB-04 also reversed established disease, produced sustained therapeutic benefits after drug withdrawal, and improved cardiac function in aged mice with spontaneous HCM. Bulk RNA sequencing and ATAC-seq demonstrated partial restoration of disease-associated transcriptional programs and chromatin accessibility. Proteomic analyses identified PHF2 (KDM7C) as a candidate target of JIB-04 in both mouse and human HCM hearts. PHF2 knockdown suppressed hypertrophic, inflammatory, and fibrotic gene expression in cardiomyocytes, macrophages, and fibroblasts, respectively. Human HCM hearts exhibited increased expression of multiple JIB-04-sensitive KDMs, including PHF2. In MYH7 R403Q induced pluripotent stem cell– derived cardiomyocytes, JIB-04 normalized disease-associated gene expression, restored connexin-43 membrane localization, and improved mitochondrial respiration. Although prolonged treatment induced reversible hepatomegaly with hepatic lipid accumulation, co-administration of the antioxidant N-acetylcysteine mitigated liver toxicity while preserving the therapeutic efficacy of JIB-04.

**Conclusions:** Pharmacological KDM inhibition prevents and reverses HCM through epigenetic remodeling of disease-associated transcriptional and chromatin programs. These findings identify KDM inhibition as a promising therapeutic strategy for HCM, establish PHF2 as a candidate mediator of disease pathogenesis, and support further development of KDM-targeted therapies.

## Introduction

Hypertrophic cardiomyopathy (HCM) is the most common inherited heart muscle disease and is primarily caused by pathogenic variants (PVs) in genes encoding sarcomeric proteins. Epidemiological studies estimate that approximately 20 million people worldwide are affected by HCM. Although many patients remain asymptomatic for years, those harboring a pathological sarcomere variants are at increased risk for progressive cardiac remodeling, heart failure, atrial fibrillation, and sudden cardiac death (SCD) ^1^. Importantly, SCD can occur in genotype-positive, phenotype-negative individuals before cardiac hypertrophy becomes detectable by echocardiography, underscoring the need for therapies that prevent disease progression rather than simply treating established structural abnormalities.

Among the most extensively studied HCM genes is *MYH7*, which encodes β-myosin heavy chain, the principal force-generating motor protein of the adult cardiac sarcomere.^2^ Patients carrying the *MYH7* p.Arg403Gln (R403Q) pathogenic variant develop hyperdynamic contraction, impaired relaxation, progressive hypertrophy, and are at particularly high risk of early SCD. ^2,3^ Current therapies for HCM include β-blockers, calcium channel blockers, anti-arrhythmic agents, septal reduction procedures, and the recently approved cardiac myosin inhibitor (CMI) mavacamten for obstructive HCM ^4–6^. Although these treatments improve symptoms and hemodynamics, they do not directly target the fibro-inflammatory and epigenetic remodeling that drives disease progression. Moreover, the recent failure of mavacamten to meet the primary endpoint in the Phase 3 ODYSSEY-HCM trial in non-obstructive HCM suggests that sarcomere-directed therapies alone may be insufficient to fully modify disease progression. These observations highlight the need for mechanism-based therapies that target the downstream molecular and cellular pathways underlying HCM.

Epigenetic regulation plays a pivotal role in cardiovascular diseases, including pathological cardiac hypertrophy. ^7–9^ In differentiated adult cardiomyocytes, the chromatin landscape is relatively stable, maintaining the transcriptional programs required for normal cardiac function. However, pathological stress and genetic mutations can remodel chromatin accessibility and gene expression, promoting maladaptive cardiac remodeling. Targeting epigenetic regulators therefore represents a selective and potentially reversible therapeutic strategy by modulating disease-associated transcriptional programs in a context- and cell type-specific manner. This concept is particularly relevant in HCM, where epigenetic dysregulation contributes to disease initiation and progression.

One hallmark of pathological cardiac remodeling is altered histone methylation^8^. Jumonji-C domain containing histone lysine demethylase (KDM) catalyze the removal of activating or repressive histone methylation marks and thereby regulate distinct transcriptional programs. We previously demonstrated that KDM4A, KDM3A, and KDM5A regulate complementary aspects of pathological cardiac remodeling. Specifically, KDM4A and KDM3A remove the repressive H3K9me3/me2 marks, whereas KDM5A demethylates the activating H3K4me3 mark. Genetic deletion of *Kdm4a* or *Kdm3a* produced minimal effects on baseline cardiac function but markedly attenuated pressure overload-induced remodeling.^10,11^ Consistent with these findings, pharmacological inhibition of multiple KDMs using the pan-KDM inhibitor JIB-04 suppressed transverse aortic constriction (TAC)-induced left ventricular hypertrophy and fibrosis and prevented sudden cardiac death in calcineurin transgenic (CnA-Tg) mice ^11, 12^. Together, these studies suggest that coordinated activation of multiple KDMs represents a common epigenetic mechanism driving pathological cardiac remodeling.

Building on these observations, we evaluated the therapeutic efficacy and molecular mechanisms of JIB-04 in the MYH6^R403Q/+^ mouse model of HCM, which carries the murine ortholog of the human MYH7 p.R403Q PV and recapitulates key features of human disease, including hyperdynamic contraction, left ventricular hypertrophy (LVH), myocyte disarray, fibrosis, diastolic dysfunction, and SCD.^2,13–15^ We demonstrate that JIB-04 both prevents disease progression and reverses established pathological remodeling in accelerated and naturally occurring HCM. Mechanistically, JIB-04 remodels disease-associated transcriptional and chromatin accessibility programs, identifies PHF2 as a candidate target within a broader KDM regulatory network, and restores conserved molecular and functional abnormalities in human MYH7 p.R403Q induced pluripotent stem cell-derived cardiomyocytes (iPSC-CM). Finally, we define the principal hepatic toxicity associated with prolonged JIB-04 treatment and demonstrate that co-administration of N-acetylcysteine preserves therapeutic efficacy while mitigating liver toxicity. Together, these findings establish KDM inhibition as a promising disease-modifying therapeutic strategy and provide a mechanistic framework for targeting convergent epigenetic pathways underlying HCM progression.

## Methods and Materials

### Plasmids, chemicals, antibodies, cell lines, and cell transfection

Lentiviral expression plasmids encoding wild-type PHF2 (pLIX402-PHF2) and constitutively active PHF2 (pLIX402 caPHF2) were obtained from Addgene ^16^. JIB-04 was obtained from the UT Dallas Chemical Core Facility. NIH3T3 cells were purchased from the American Type Culture Collection (ATCC, Manassas, VA, USA). Anti-PHF2 antibody (clone D5Q8E, #3497, rabbit monoclonal) was purchased from Cell Signaling Technology (Danvers, MA, USA). Anti-connexin 43 antibody (ACC-201-GP) was purchased from Alomone Labs (Jerusalem, Israel). PHF2 siRNAs were purchased from Sigma-Aldrich (human: SASI_Hs02_00339519 and SASI_Hs01_00185572; rat: SASI_Rn02_00227885 and SASI_Rn02_00227886). Plasmid DNA and siRNA transfections were performed using Lipofectamine 3000 and Lipofectamine RNAiMAX (Thermo Fisher Scientific), respectively, according to the manufacturer’s instructions.

### Human tissues, mouse husbandry, and JIB-04 treatment

Human hypertrophic septal tissue samples were obtained under a protocol approved by the Institutional Review Board of the University of Texas Southwestern Medical Center (IRB #STU 082017-072). All animal experiments were approved by the Institutional Animal Care and Use Committee (IACUC) of the University of Texas Southwestern Medical Center.

Mice were maintained on standard rodent chow (Teklad Global 2916). Generation of the humanized Myh6^R403Q/+^ mice containing human analogous Myh7 R403Q pathogenic variant on a C57BL/6 background has been described previously ^13^. Experimental Myh6^R403Q/+^ mice and wild-type littermate controls were generated by crossing Myh6^R403Q/+^ mice with wild-type (Myh6^+/+^) mice. Genotyping was performed by polymerase chain reaction (PCR) using the primers listed in supplemental Table 1. To accelerate the onset of HCM, mice were fed a custom Teklad Global 2916-based chow containing cyclosporine A (1 g/kg; Alfa Aesar) and blue food dye (0.2 g/kg).

JIB-04 stock solution was prepared in dimethyl sulfoxide (DMSO) and diluted in a vehicle consisting of 10% DMSO, 10% ethoxylated castor oil (Cremophor EL), and 80% sterile water. The vehicle control group received the same formulation without JIB-04. Fresh dosing solutions were prepared on the day of treatment and administered to mice by oral gavage.

### Transthoracic echocardiography

Cardiac function was evaluated by two-dimensional transthoracic echocardiography using a Vevo 2100 high-resolution imaging system equipped with an MS400 transducer (VisualSonics). Systolic function was assessed in conscious mice using M-mode images acquired from the parasternal short-axis view at the level of the papillary muscles. Measured parameters included LVEF, FS, LVIDd, LVIDs, IVSd, and LVPWd. FS and LVEF were calculated using Vevo software (VisualSonics).

Diastolic function was assessed in anesthetized mice from the apical four-chamber view using pulsed-wave Doppler and tissue Doppler imaging at the level of the mitral valve. Anesthesia was induced with 5% isoflurane in oxygen and maintained at 1.0–1.5% isoflurane in oxygen throughout imaging acquisition. Body temperature was maintained at 37°C using a heated platform and heart rate was continuously monitored and maintained between 415 and 460 beats per minute (bpm). Diastolic parameters included peak early (E) and late (A) mitral flow velocities, the E/A ratio, early diastolic mitral annular tissue velocity (e′), the E/e′ ratio, isovolumic relaxation time (IVRT), and deceleration time (DT). All measurements were averaged from at least three consecutive cardiac cycles and were obtained by an experienced operator blinded to the experimental groups.

### Histology

Mouse hearts were excised and immersed in cardioplegic PBS containing 0.2 M KCl for 5 min before fixation in 4% paraformaldehyde in PBS overnight. Tissues were then dehydrated in 70% ethanol, processed, and embedded in paraffin. Serial transverse sections were cut at 500-μm intervals, mounted on glass slides, and stained with hematoxylin and eosin (H&E), picrosirius red, or Masson’s trichrome. Images were captured on a BZ-X800 all-in-one microscope (Keyence) at ×10 or ×40 magnification. Analyses were performed with ImageJ (NH) and Adobe Photoshop.

### Demethylase assay

Nuclear extracts were isolated from mouse heart tissue using the EpiQuik Nuclear Extraction Kit (Epigentek, OP-0002-1) according to the manufacturer’s instructions. Nuclear extracts were used as the enzyme source to measure demethylase activity against exogenous biotinylated H3K4me3, H3K9me3, or H3K27me3 histone peptide substrates, as previously described ^17–19^. H3K4me3 demethylation was measured using the Epigenase JARID Demethylase Activity/Inhibition Assay Kit (Epigentek, P-3083), whereas H3K9me3 demethylation was measured using the Epigenase JMJD2 Demethylase Activity/Inhibition Assay Kit (Epigentek, P-3081).

Reactions were carried out in a final volume of 50 μL containing 1 mM α-ketoglutarate, 2 mM ascorbate, 100 μM FeSO₄·7H₂O (Sigma, F7002), 50 ng biotinylated histone peptide substrate (H3K4me3, H3K9me3, or H3K27me3), 0.25× EDTA-free protease inhibitor (Roche, 05 892 791 001), and 12 μg nuclear extract. Reaction mixtures were incubated at 37°C for 2 h. Data are presented as arbitrary fluorescence units (AU) measured by the assay.

### RNA isolation, RNA sequencing, ATAC-sequencing, and heatmap generation

Total RNA was isolated from heart tissue using TRIzol reagent according to the manufacturer’s instructions. Bulk RNA sequencing was performed by Admiral Health (Boston, MA). Raw paired-end RNA-seq FASTQ files were assessed for quality using FastQC v0.12.1^20^ and FastQ Screen v0.15.3^21^. Reads were aligned to the mouse reference genome (mm10) with STAR v2.5.3a ^22^. Gene-level counts were quantified using featureCounts^23^, differential expression analysis was performed using edgeR ^24^, and Pathway enrichment was performed by gene set enrichment analysis (GSEA) using the WebGestaltR package ^25^.

Nuclei were isolated from heart tissues following published protocol ^26^. ATAC-seq libraries were prepared using ATAC-seq kit (Active motif,#53150) according to manufacturer’s instruction. Sequencing was performed by Admiral Health (Boston, MA). Raw paired-end ATAC-seq reads were assessed for quality using FastQC v0.12.1 ^20^ and FastQ Screen v0.15.3 ^21^. Adapter sequences and low-quality bases were removed using fastp v0.24.0 ^27^. Trimmed reads were aligned to the mouse reference genome (mm10) using Bowtie2 v2.5.4 ^28^. Final BAM files were generated using Sambamba v1.0.1 ^29^ and deepTools alignmentSieve v3.5.5 ^30^ by removing mitochondrial reads and PCR duplicates, retaining properly paired reads with mapping quality ≥10, and excluding secondary, supplementary, QC-failed, and ENCODE mm10 blacklist v2 ^31^ Accessible chromatin regions were identified using MACS3 HMMRATAC v3.0.3 ^32^. Peak annotation was performed using ChIPseeker with mm10 gene models ^33^. Normalized BigWig files were generated using deepTools v3.5.5 ^30^ with RPGC normalization. Aggregate signal profiles were generated over the ENCODE Registry of candidate cis-regulatory elements (cCREs) ^34^ for mm10 using deepTools v3.5.5.

For heatmap visualization, differentially expressed genes (DEGs) associated with pathways of interest were identified from the RNA-seq analysis. Genes were ranked according to their average normalized expression across all samples, and representative DEGs of interests within each pathway were selected for visualization. The corresponding normalized expression matrix was exported and uploaded to the Bioinformatics Platform Heatmap Tool to generate hierarchically clustered heatmaps illustrating the expression patterns of the selected genes^35^.

### Biotin-JIB-04 pull-down and proteomic analysis

Nuclear extracts (500 μg total protein) were prepared from diseased mouse and human HCM hearts using nuclear lysis buffer (25 mM Tris-HCl, pH 8.0, 150 mM NaCl, 2 mM MgCl₂, 0.5% NP-40, supplemented with protease and phosphatase inhibitor cocktails). Biotin-JIB-04 (3 μL of a 100 μM stock) or biotin alone (negative control) was first immobilized on streptavidin-coated beads and then incubated with the nuclear extracts in the presence of 1mM TCEP, 500 μM α-ketoglutarate, 10 μM ferrous iron (Fe²⁺), and 1 mM ascorbic acid to maintain KDM catalytic activity. The reactions were incubated at 4°C for 2 h with gentle rotation.

After incubation, the beads were washed three times with PBS containing 0.1% NP-40 to remove nonspecifically bound proteins. Bound proteins were resolved by SDS-PAGE and visualized by Coomassie Brilliant Blue staining. Following destaining (40% methanol, 10% acetic acid) and rinsing with distilled water, each gel lane was excised into approximately 1 × 1 mm pieces and subjected to in-gel tryptic digestion followed by liquid chromatography-tandem mass spectrometry (LC-MS/MS) for protein identification at the UT Southwestern Proteomics Core Facility. MS data were analyzed using Proteome Discoverer 3.0 (Thermo Fisher Scientific) and searched against the UniProt mouse or human protein database, as appropriate.

### Seahorse extracellular flux analysis

Mitochondrial oxygen consumption rate (OCR) was measured in live cardiomyocytes using a Seahorse XF Analyzer. Sequential injections of oligomycin, FCCP, and rotenone/antimycin A were used to assess ATP-linked respiration, maximal respiratory capacity, and non-mitochondrial respiration, respectively. Extracellular acidification rate (ECAR) was measured in parallel as an indicator of glycolytic activity.

### Statistical analysis

All data are presented as mean ± SEM or mean ± SD, as specified in each figure legend, along with the number of biological replicates (n). Unless otherwise noted, each analysis was conducted across at least three independent experiments, with 2-3 biological samples per experimental group. Before running any comparisons, we tested each dataset for normality using the Shapiro-Wilk test in GraphPad Prism 10, applying a significance threshold of 0.05. For two-group comparisons, we used an unpaired two-tailed Student’s t-test when data were normally distributed, or the Mann-Whitney test when they were not. When comparing three or more groups, we applied one-way ANOVA followed by Tukey’s post hoc test to account for multiple comparisons. All statistical analyses were carried out in GraphPad Prism 10, and differences were considered statistically significant at P < 0.05.

## Results

### 1. JIB-04 prevents disease progression in the Myh6^R403Q/+^ mouse model of HCM

*Myh6^R403Q/+^* mice develop spontaneous HCM with age.^2,13–15^ To accelerate disease progression, mice were fed a 0.1% cyclosporine A (CsA) chow diet beginning at 10 weeks of age. This model recapitulates key features of HCM by exacerbating mitochondrial stress and calcium-calcineurin signaling in the setting of the sarcomeric mutation. ^2,13–15^ JIB-04 treatment (30 mg/kg by oral gavage, three times per week) was initiated concurrently with CsA and continued for 10 weeks (Fig. 1A).

**Figure 1.**
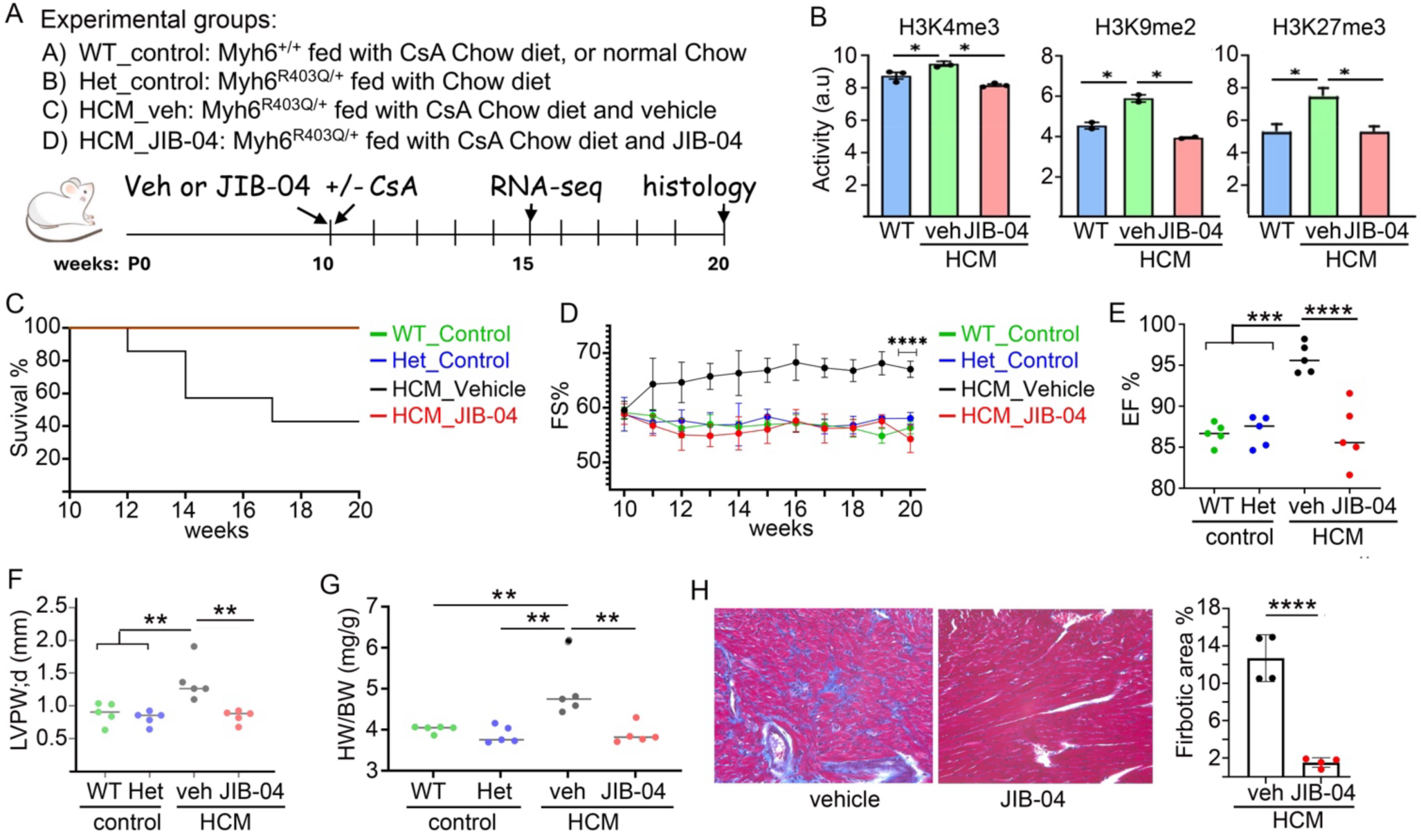
JIB-04 prevents HCM progression in Myh6^R403Q/+^ mice. (A) Experimental design and study groups. Four groups were analyzed throughout this study: WT_control (WT littermates on CsA diet), Het_control (*Myh6^R403Q/+^* on standard chow), HCM_veh (*Myh6^R403Q/+^* mice fed a 0.1% CsA diet and treated with vehicle), and HCM_JIB-04 (*Myh6^R403Q/+^* mice fed a 0.1% CsA diet and treated with JIB-04),. Ten-week-old mice were fed with either CsA-containg or control diet and treated with JIB-04 (30mg/kg) or vehicle by oral gavage three time a week for 10 weeks. (B) Demethylase activities in hearts from the indicated groups after 10-weeks treatments (n=3 per group). (C) Kaplan-Meier survival analysis (n=7 per group). (D) Fractional Shortening (FS%). (E-H) Echocardiographic and histological measurements obtained at the end of the 10-week treatment period, including Lleft ventricular ejection fraction (EF%) (E), Left ventricular posterior wall thickness at end-diastole (LVPW;d) (F), heart weight/body weight ratio (HW/BW) (G), representative trichrome staining (H, left) and quantification of myocardial fibrosis (H, right). Data are mean ± SD. *\*, p < 0.05; **, p < 0.01; ***, p < 0.005, ****, p<0.001*.

JIB-04 significantly reduced the elevated H3K9me3, H3K4me3, and H3K27me3 demethylase activities observed in HCM hearts (Fig. 1B), confirming target engagement of JIB-04 in vivo. Treatment completely prevented sudden cardiac death (SCD) (Fig. 1C), preserved fractional shortening (FS) and ejection fraction (EF) at levels comparable to wild-type controls (Figs. 1D and 1E), and attenuated pathological remodeling, as evidenced by reduced left ventricular posterior wall thickness at end-diastole (LVPW;d), lower heart weight (HW)-to-body weight (BW) ratios, and markedly decreased myocardial fibrosis (Figs. 1F–1H). Consistent with our previous findings,^11^ JIB-04 had no detectable effects on cardiac structure or function in wild-type mice, supporting selective therapeutic activity in diseased hearts.

### 2. JIB-04 reverses established HCM

*Myh6^R403Q/+^* mice were randomized to receive JIB-04 or vehicle beginning 5 weeks after HCM induction by CsA diet (Fig. 2A), when hypercontractility was already evident (Fig. 1D). After 5 weeks of treatment, a subset of mice discontinued JIB-04 while continuing the CsA diet to assess the durability of therapeutic response. JIB-04 normalized cardiac function by reversing hypercontractility (Fig. 2B), reduced cardiac hypetrophy and HW/BW ratios (Fig. 2C), and markedly attneuated established myocardial fibrosis (Fig. 2D). Notably, these functional improvement were maitained for at least 5 weeks after drug withdrawal despite continued CsA exposure (Fig. 2B), suggesting sustained disease modification rather than transient pharmacologic suppression.

**Figure 2.**
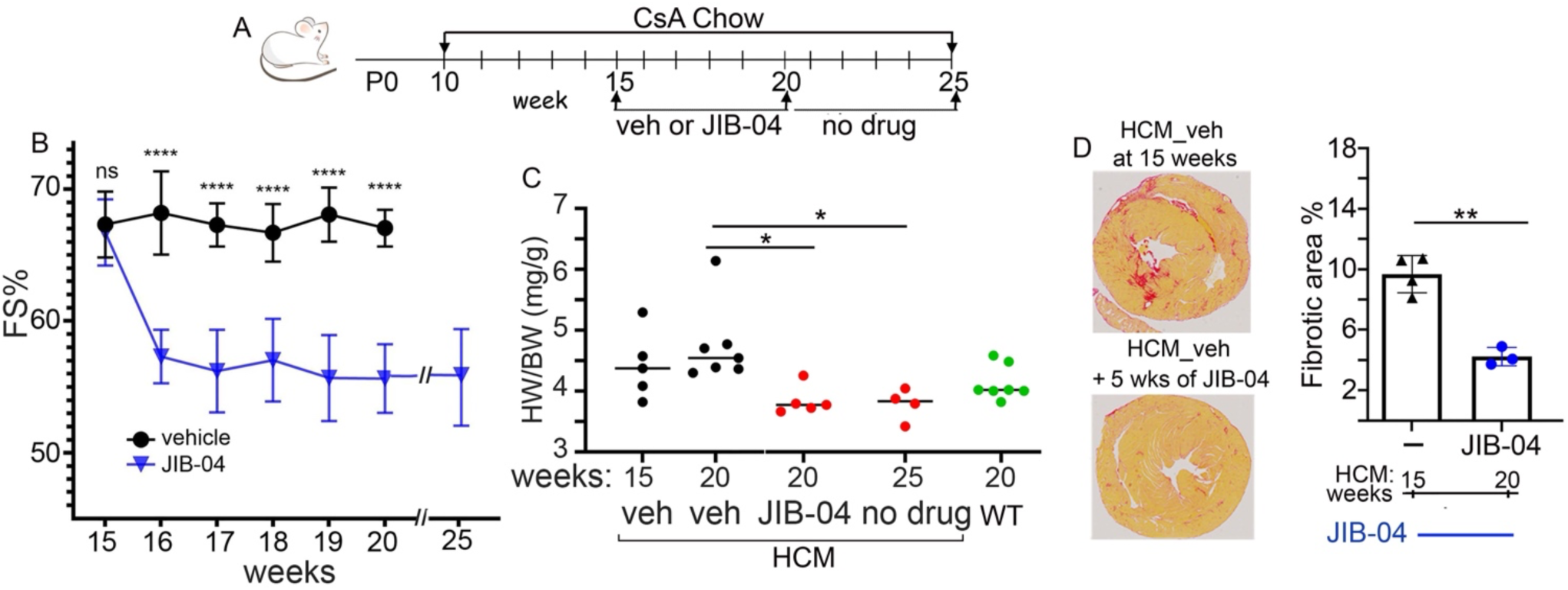
JIB-04 reverses established HCM. (A) Experimental design. *Myh6^R403Q/+^* mice were fed a 0.1% CsA diet beginning at 10 weeks of age. After 5 weeks of CsA-induced HCM, mice were randomized to receive vehicle or JIB-04 (30 mg/kg by oral gavage, 3 times per week) for 5 weeks while remaining on the CsA diet. A subset of JIB-04-treated mice was followed for an additional 5 weeks after drug withdrawal. (B) Serial measurements of FS% in vehicle- and JIB-04-treated HCM mice. (C) HW/BW in HCM mice at treatment initiation (15 weeks), after 5 weeks of vehicle or JIB-04 treatment (20 weeks), and after a 5-week drug withdrawal period (25 weeks). Age-matched wild-type (WT) mice are shown for comparison. (D) Left: Representative Picrosirius red-stained heart section from HCM mice at treatment initiation (15 weeks) and after 5 weeks of JIB-04 treatment (20 weeks). Right: Quantification of fibrotic areas in the corresponding HCM hearts. Data are presented as mean ± SD, *, p < 0.05; **, p < 0.01; ****. p < 0.001.

### 3. JIB-04 improves cardiac function in aged HCM mice

To determine whether JIB-04 is effective in naturally progressing disease, we evaluated its effect in aged *Myh6^R403Q/+^* mice that spontaneously develop HCM without CsA treatment. Unlike the accelerated CsA-induced model, disease progression in aged mice is heterogeneous, with some animals exhibiting established diastolic dysfunction while others retain normal diastolic function at treatment initiation. Despite this variability, JIB-04 attenuated hypercontractility, a hallmar of this model, as reflected by reduced fractional shortening (%FS) (Fig 3A), Vehicle-treated HCM mice exhibited progressive worsening of diastolic function over the 5-week study period, as indicated by an increase in E/e′ ratio, whereas JIB-04 prevented this deterioration and reduced E/e′ in mice with elevated baseline values, resulting in significantly improved diastolic function compared with vehicle-treated HCM mice (Fig. 3B). JIB-04 also reduced left atrial size to levels comparable to those of age-matched wild-type controls (Fig. 3C) and inhibited fibrosis (Fig. 3D). These findings demonstrate that the therapeutic benefits of JIB-04 extend beyond the accelerated CsA-induced model and support its efficacy in naturally progressive HCM.

**Figure 3.**
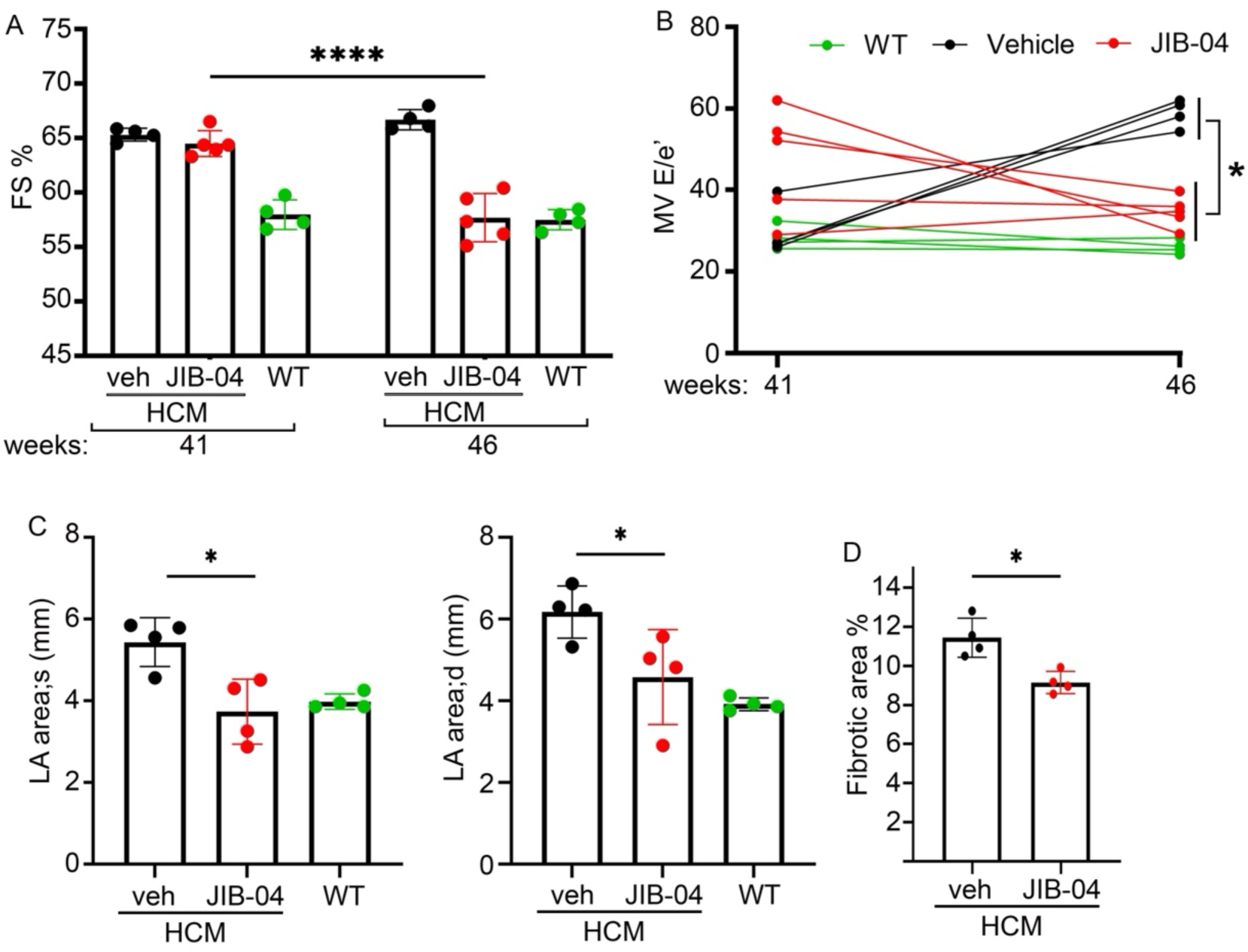
JIB-04 improves cardiac function in aged HCM mice. Forty-one-week-old Myh6^R403Q/+^ mice were treated with vehicle or JIB-04 for 5 weeks. Cardiac structure and function were assessed by echocardiography before and after treatment. (A) Fractional shortening (FS%). (B) Individual changes in E/e′ ratio in WT mice (green), vehicle-treated HCM mice (black), and JIB-04-treated HCM mice (red). (C) Left atrial area at systole and diastole after 5 weeks of treatment (46 weeks). (D) Fibrotic area % of aged Myh6^R403Q/+^ (46 weeks) mice after 5 weeks treatment with vehicle or JIB-04. Data are presented as mean ± SD. *, p < 0.05; ****, p < 0.001.

### 4. JIB-04 reverses disease-associated transcriptional programs in HCM

We performed bulk RNA-seq on *Myh6^R403Q/+^* hearts after 5 weeks of HCM induction, comparing JIB-04-treated (HCM_JIB-04) and vehicle-treated (HCM_veh) mice together with non-diseased *Myh6^R403Q/+^* (Het_control) and wild-type (WT_control) controls (Fig. 1A). Differential gene expression analysis revealed extensive transcriptional remodeling in diseased HCM hearts, a substantial proportion of which was reversed by JIB-04 treatment (Figs. 4A and 4B). Specifically, JIB-04 suppressed 38% of genes upregulated in HCM and restored 36% of genes downregulated in HCM toward control levels (Fig. 4B).

**Figure 4.**
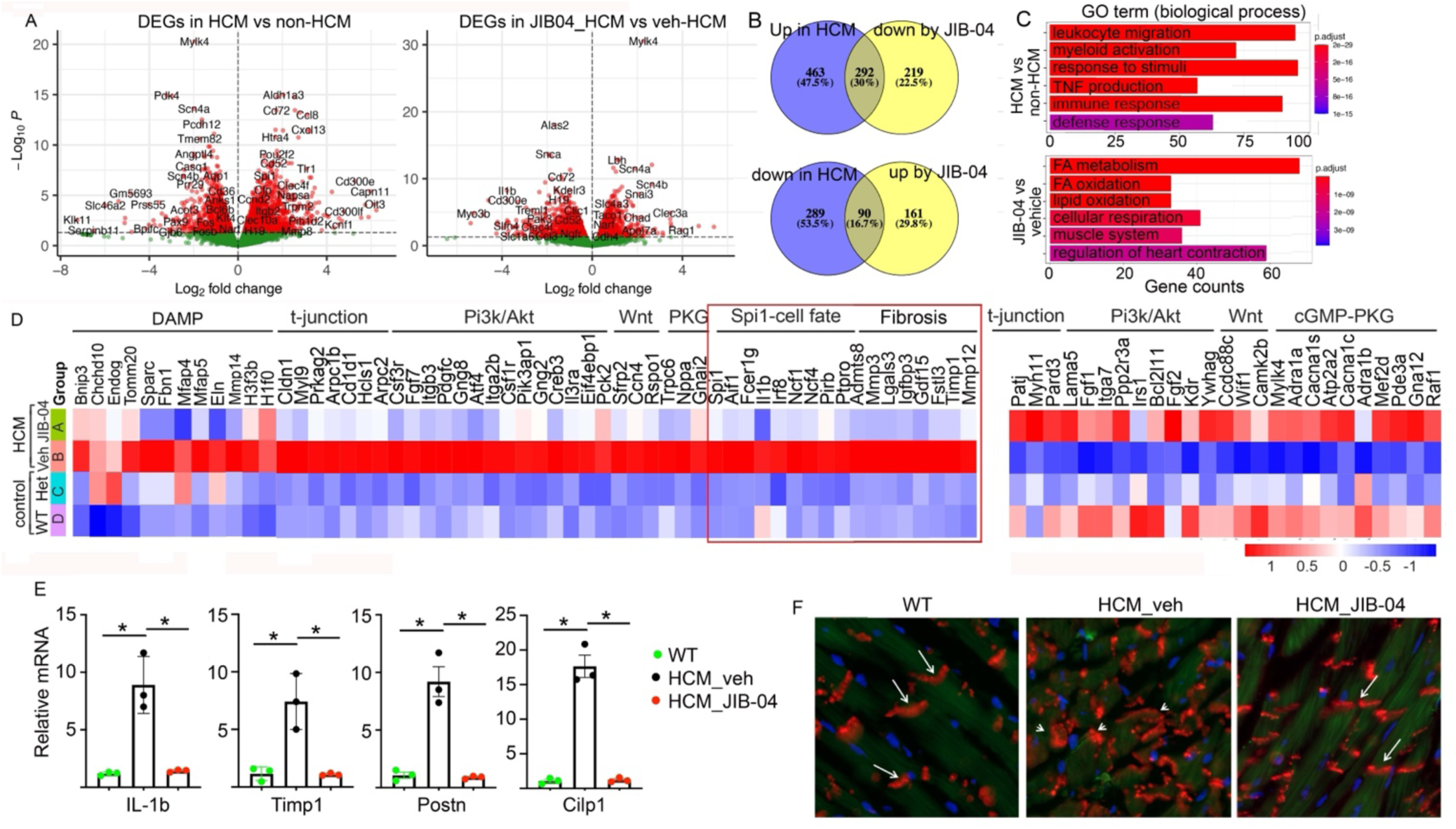
JIB-04 remodels disease-associated transcriptional programs in HCM. (A) Volcano plots showing differentially expressed genes (DEGs) in HCM_veh versus non-HCM controls (left) and HCM_JIB-04 versus HCM_veh hearts (right). Each dot represents a single gene. Red dots indicate significantly upregulated or downregulated DEGs that meet the predefined statistical significance threshold, whereas green dots represent genes that do not meet the significance threshold. (B) Venn diagrams illustrating overlap between genes dysregulated in HCM and genes normalized by JIB-04 treatment. Upper panel, genes upregulated in HCM and downregulated by JIB-04; lower panel, genes downregulated in HCM and upregulated by JIB-04. (C) Gene Ontology (GO) biological process enrichment analysis of genes increased in HCM relative to non-HCM controls (top) and genes increased by JIB-04 treatment relative to vehicle-treated HCM hearts (bottom). (D) Heat maps showing representative genes from selected pathways altered in HCM and restored by JIB-04, including DAMP signaling, t-tubule/junctional pathways, PI3K/AKT signaling, Wnt signaling, PKG signaling, Spi1-associated cell fate, and fibrosis programs. (E) Relative expression of genes involved in inflammation and fibrosis in wild-tpe and HCM heart treated with vehicle or JIB-04. Data are presented as mean ± SD. *, P < 0.05. (F) Representative immunostaining for connexin-43 (Cx43, red), cardiomyocytes autofluoresence (green), and nuclei (DAPI, blue) in WT, HCM_veh, and HCM_JIB-04 hearts. Arrows indicate Cx43 localized at intercalated discs in WT and JIB-04 treated hearts. Arrowheads in HCM_veh hearts indicate lateralized and intracellular Cx43. JIB-04 restored Cx43 localization toward the WT pattern.

Gene ontology (GO) analysis demonstrated enrichment of immune activation, inflammatory signaling, and stress-response pathways in diseased HCM hearts, whereas genes normalized by JIB-04 were enriched in pathways related to cardiac contraction, muscle function, lipid metabolism, and fatty acid oxidation (Fig. 4C). Consistent with these findings, pathway analysis showed normalization of multiple disease-associated signaling programs, including damage-associated molecular pattern (DAMP) signaling, T-tubule and cell-cell junctional organization, PI3K-AKT, Wnt, cGMP-PKG, Spi1-associated inflammatory, and fibrosis pathways (Fig. 4D). Selected inflammatory and fibrotic genes identified by RNA-seq were further validated by qRT-PCR (Fig. 4E). Immunofluorescence staining of the gap junction protein connexin-43 (Cx43) revealed disrupted intercellular localization, with increased intracellular and lateral Cx43 staining in diseased HCM hearts compared with WT controls. JIB-04 partially restored Cx43 localization toward the normal membrane-associated pattern (Fig. 4F). Collectively, these findings indicate that JIB-04 coordinately restores transcriptional programs governing myocardial function, metabolism, inflammation, and cellular signaling in HCM.

### 5. PHF2 (KDM7C) is a candidate JIB-04 target in HCM

To identify candidate cardiac targets of JIB-04, we performed biotin-JIB-04 pulldown followed by mass spectrometry using nuclear lysates from mouse and human HCM hearts. The human samples represented diverse genetic etiologies. PHF2 (KDM7C) was identified in the pulldown from both human and mouse hearts (Fig. 5A; Supplemental Table 2). The PKA catalytic subunit PRKACA, and PKA associated proteins AKAP13 and PRKAR2A were also identified among the proteins shared between species. In addition, several chromatin-associated proteins, RBBP4, SMARCC2 and ACTL6A were recovered in the pulldowns (Supplemental Table 2). Together, these findings nominate PHF2 and components of a PKA-associated chromatin-regulatory complex as candidate JIB-04-interacting proteins.

**Figure 5.**
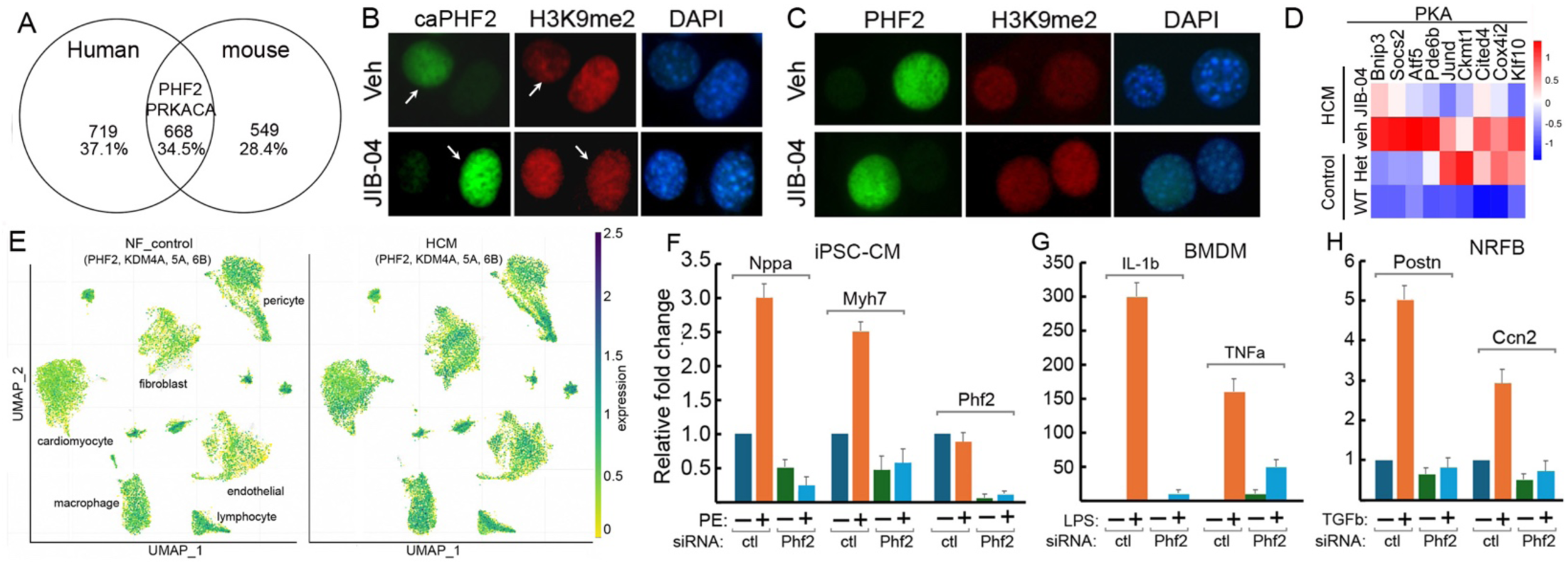
PHF2 is a candidate target of JIB-04 in HCM. Biotin-labeled JIB-04 were incubated with nuclear lysates from mouse and human HCM hearts and precipitated using streptavidin beads. Pull-down proteins were identified by mass-spectrometry analysis. (A) Shared proteins identified by biotin-JIB-04 pulldown from mouse and human HCM hearts. (B-C) Immunofluorescence staining of PHF2, H3K9me2, and DAPI in NIH3T3 cells expressing constitutively active PHF2 (caPHF2; B) or non-activated PHF2 (C) and treated with vehicle or JIB-04. (D) Heatmap of PKA-responsive genes in the indicated mouse groups. (E) Aggregate expression of PHF2, KDM4A, KDM5A, and KDM6B in human non-failing (NF) and HCM hearts. (F-H) PHF2 knockdown attenuated hypertrophic, inflammatory, and profibrotic gene expression programs in iPSC-CMs, BMDMs, and NRFBs stimulated with PE, LPS, or TGFβ, respectively. Data are presented as mean ± SEM; n = 3 independent experiments.

PHF2 is activated by PKA-dependent phosphorylation and catalyzes demethylation of H3K9me2. ^16,36^ Consistent with this mechanism, overexpression of constitutively active PHF2 (caPHF2) in NIH3T3 cells reduced H3K9me2 levels, whereas JIB-04 blocked this effect (Fig. 5B). In contrast, wild-type PHF2 produced minimal changes in H3K9me2 and was largely insensitive to JIB-04 (Fig. 5C). Moreover, canonical PKA-responsive genes were upregulated in *Myh6^R403Q/+^* hearts and further induced following HCM induction (Fig. 5D), consistent with increased β-adrenergic/PKA signaling in HCM. ^37,38^ These findings suggest PHF2 as a candidate functional target of JIB-04 and raise the possibility that JIB-04 modulates a PKA–PHF2 signaling axis during HCM progression.

To evaluate the clinical relevance of these findings, we analyzed a published human HCM single-nuclei RNA-sequencing dataset ^39^ and identified coordinated upregulation of a KDM module comprising PHF2, KDM4A, KDM5A, and KDM6B in genetically heterogeneous HCM hearts (Fig. 5E). We next examined the functional role of PHF2 in major cardiac cell populations. PHF2 knockdown in human induced pluripotent stem cell-derived cardiomyocytes (iPSC-CMs), mouse bone marrow-derived macrophages (BMDMs), and neonatal rat fibroblasts (NRFBs), significantly attenuated phenylephrine-, lipopolysaccharide-, and TGFβ-induced pathological gene expression, respectively (Figs. 5F-5H). Consistent with these observations, RNA-seq demonstrated that JIB-04 suppressed many of the same hypertrophic, inflammatory, and profibrotic transcriptional programs. Collectively, these findings identify PHF2 as a candidate mediator of JIB-04 activity and support a model in which JIB-04 regulates pathological remodeling through a broader KDM network operating across multiple cardiac cell types.

### 6. JIB-04 reverses disease-associated phenotypes in human HCM iPSC-CM

To evaluate the translational relevance of JIB-04, we examined its effects in patient-derived iPSC-CM carrying the MYH7 R403Q mutation (HCM1) and an isogenic CRISPR-corrected control line (HCM1^corr^).^13^ Several genes dysregulated in *Myh6^R403Q/+^*hearts and restored by JIB-04 exhibited similar expression changes in HCM1 relative to HCM1^corr^ and were likewise normalized by JIB-04 treatment (Fig. 6A). JIB-04 also restored membrane localization of Cx43, which was disrupted in HCM1 but preserved in HCM1^corr^ (Fig. 6B).

**Figure 6.**
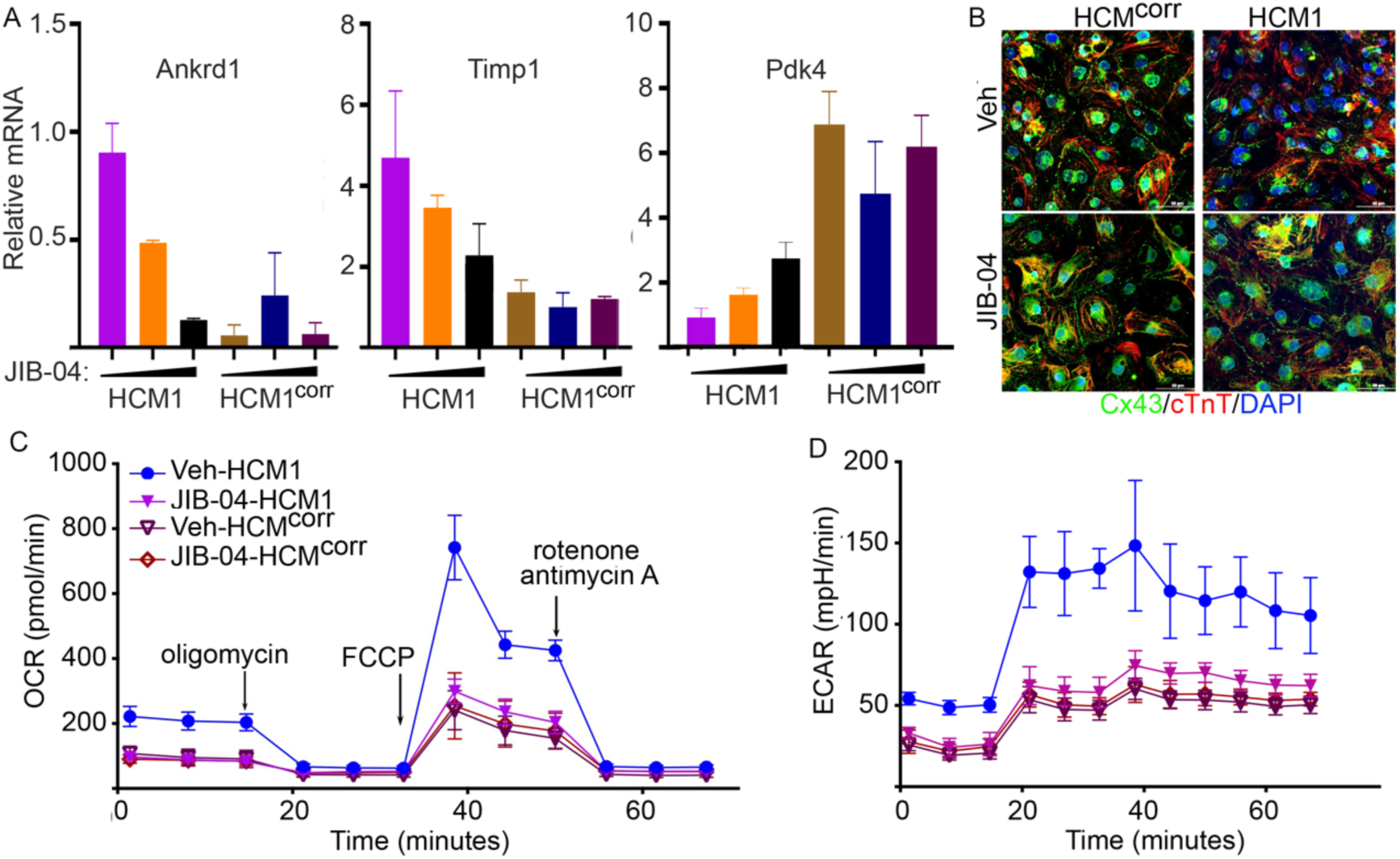
JIB-04 reverses disease-associated phenotypes in human HCM iPSC-derived cardiomyocytes. Human MYH7 R403Q HCM iPSC-derived cardiomyocytes (HCM1) and isogenic gene-corrected control cardiomyocytes (HCM1^corr^) were treated with vehicle or JIB-04 and analyzed by molecular, structural, and metabolic assays. (A) Expression of the hypertrophic marker Ankrd1, fibrotic/injury marker Timp1, and metabolic regulator Pdk4 measured by qRT-PCR. (B) Immunofluorescence staining for connexin-43 (Cx43, green), cardiac troponin T (cTnT, red), and nuclei (DAPI, blue). (C-D) Seahorse extracellular flux analysis showing oxygen consumption rate (OCR; C) and extracellular acidification rate (ECAR; D). Oligomycin, FCCP, and rotenone/antimycin A were sequentially injected to assess ATP-linked respiration, maximal respiratory capacity, and non-mitochondrial respiration, respectively. JIB-04 normalized disease-associated gene expression, restored membrane localization of Cx43, and improved mitochondrial function in HCM1 cardiomyocytes with minimal effects in isogenic corrected controls. Data are presented as mean ± SD. n = 3 independent experiments for qRT-PCR and n = 10 wells per group for Seahorse analyses.

We next measured oxygen consumption rate (OCR) in live HCM1 cells using a Seahorse XF Analyzer. HCM1 exhibited elevated basal respiration and a markedly exaggerated FCCP-stimulated maximal respiratory capacity compared with HCM^corr^, indicative of mitochondrial hyperactivation. Treatment with JIB-04 significantly reduced both basal and maximal OCR in HCM1 cells, restoring a respiratory profile comparable to HCM^corr^ cells (Fig. 6C). Extracellular acidification rate (ECAR) was measured in parallel as an index of glycolytic flux. HCM1 cardiomyocytes displayed increased ECAR, particularly under metabolic challenge, consistent with enhanced reliance on glycolysis. In contrast, HCM^corr^ cardiomyocytes maintained lower and more stable ECAR levels. JIB-04 treatment attenuated glycolytic activation in HCM1 cells without inducing compensatory glycolytic upregulation, indicating improved metabolic efficiency rather than energetic suppression (Fig. 6D).

### 7. JIB-04 partially restores chromatin accessibility in HCM

To determine whether JIB-04 remodels the epigenetic landscape in HCM, we performed ATAC-seq on wild-type (WT_control), non-diseased *Myh6^R403Q/+^* (Het_control), diseased HCM (HCM_veh), and JIB-04-treated HCM (HCM_JIB-04) hearts. Compared with WT_control hearts, both Het_control and HCM_veh hearts exhibited reduced promoter accessibility together with increased accessibility at distal intergenic regions (Fig. 7A). The highly similar accessibility profiles of Het_control and HCM_veh hearts suggest that substantial chromatin remodeling occurs before overt disease develops and persists throughout disease progression. JIB-04 treatment partially restored chromatin accessibility toward the WT pattern.

**Figure 7.**
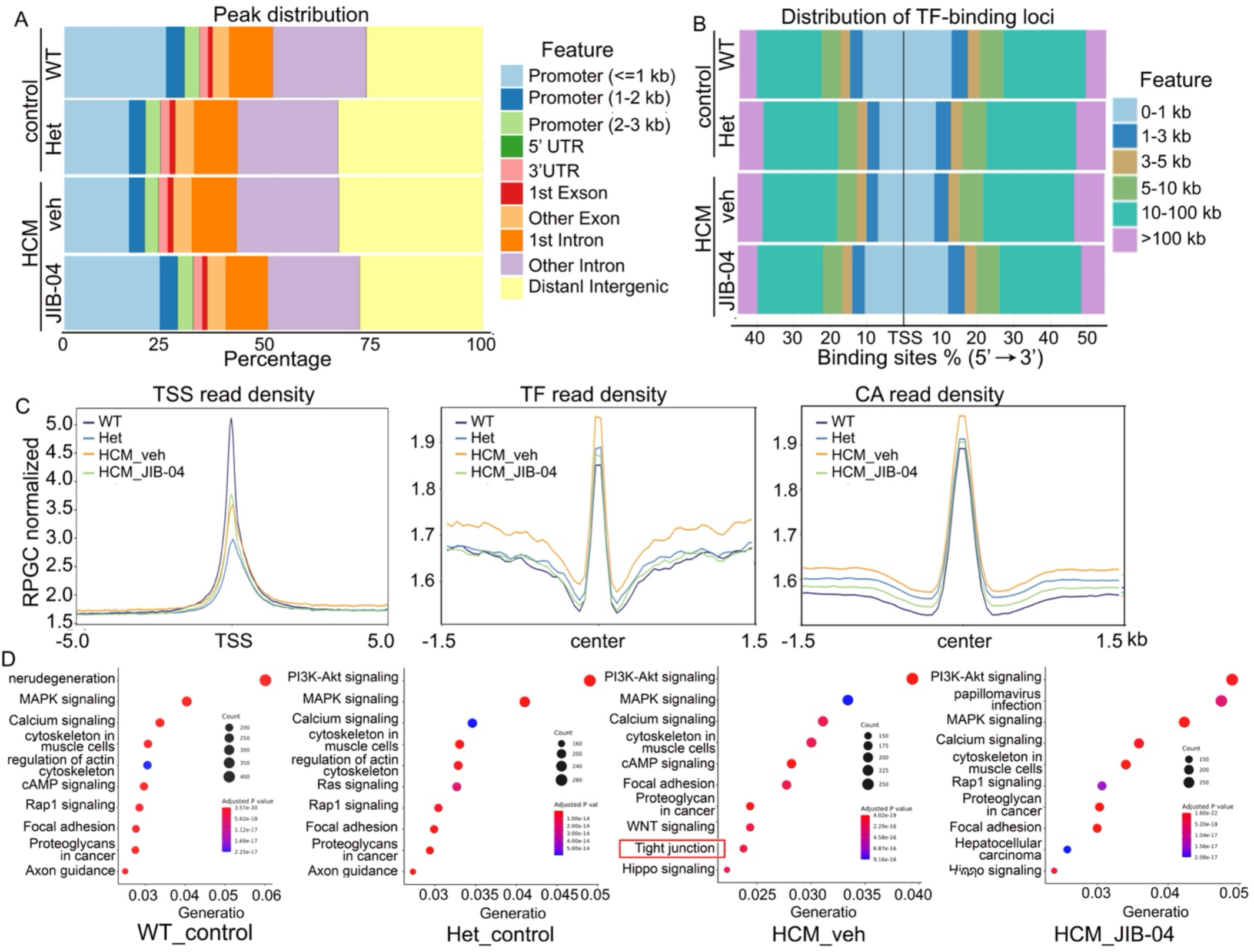
JIB-04 remodels chromatin accessibility in HCM. (A) Distribution of ATAC-seq peaks across genomic features in WT_control, Het_control, HCM_veh, and HCM_JIB-04 hearts. Genomic annotations include promoter, untranslated region (UTR), exon, intron, and distal intergenic regions. (B) Distribution of transcription factor (TF)-binding loci. Peaks were categorized according to their distance from transcription start sites (TSSs) and plotted as percentages of total binding sites. Het_control hearts displayed reduced promoter accessibility and increased distal intergenic accessibility compared with WT controls, abnormalities that persisted in HCM_veh hearts and were partially normalized by JIB-04. (C) Aggregate ATAC-seq signal profiles showing normalized read density around transcription start sites (TSS; left), transcription factor-binding sites (TF; middle), and accessible chromatin regions (CA; right). HCM hearts exhibited reduced and broadened TSS accessibility compared with WT controls, whereas JIB-04 partially restored accessibility toward the WT pattern. (D) KEGG pathway enrichment analysis of genes associated with accessible chromatin regions in WT_control, Het_control, HCM_veh, and HCM_JIB-04 hearts. Bubble size represents the number of genes associated with each pathway, and color indicates the adjusted P value. PI3K-AKT signaling, MAPK signaling, calcium signaling, cytoskeletal organization, and focal adhesion were among the top enriched pathways across the four groups. Tight junction signaling appeared only in the HCM_veh dataset.

Consistent with these findings, analysis of chromatin accessibility surrounding predicted transcription factor (TF)-binding sites revealed altered accessibility patterns in mutant hearts (Fig. 7B). Compared with WT controls, Het_control and HCM_veh hearts exhibited reduced accessibility in promoter-proximal regions surrounding transcription start sites (TSSs), accompanied by a relatively increase in accessibility at more distal regulatory regions. JIB-04 partially normalized this distribution, suggesting restoration of chromatin accessibility toward the WT pattern.

Genome-wide accessibility profiles further supported these observations. TSS read density was reduced and broadened in both Het_control and HCM_veh hearts relative to WT controls (Fig. 7C), indicating diminished promoter accessibility. The highly similar TSS profiles observed in these two groups suggest that altered promoter accessibility is an early consequence of the Myh6 R403Q mutation that precedes overt HCM. In contrast, TF-binding sites and accessible chromatin (CA) rad-density profiles showed elevated flanking accessibility in HCM_veh hearts compared with WT and Het_control hearts, whereas JIB-04 shifted both profiles toward the WT pattern. Together, these findings indicate that the Myh6 R403Q mutation induces early epigenetic remodeling that persists during disease progression and is partially reversed by KDM inhibition.

To gain insight into the biological pathways associated with accessible chromatin regions, we performed pathway enrichment analysis of ATAC-seq peaks (Fig. 7D). Across all groups, accessible regions were enriched for signaling pathways implicated in cardiac remodeling, including PI3K-Akt, MAPK, calcium signaling, cAMP signaling, Rap1 signaling, focal adhesion, and regulation of the actin cytoskeleton. Notably, tight junction signaling was uniquely enriched in HCM_veh hearts, whereas PI3K-Akt signaling was already differentially represented in Het_control hearts, supporting the presence of early chromatin remodeling before overt disease onset. Following JIB-04 treatment, the accessibility landscape shifted toward pathways more similar to WT hearts. Importantly, many pathways identified by ATAC-seq overlapped with those altered in the bulk RNA-seq dataset (Fig. 4), suggesting that changes in chromatin accessibility contribute to the transcriptional reprogramming observed during HCM progression and its reversal by JIB-04.

Together, these results identify chromatin remodeling as an early consequence of the R403Q mutation that precedes overt HCM and demonstrate that JIB-04 partially reverses disease-associated epigenomic alterations. The presence of chromatin accessibility changes in Het_control hearts indicates that epigenetic dysregulation occurs before detectable pathological remodeling, supporting the concept that chromatin remodeling is an early pathogenic event in HCM and is therapeutically reversible by KDM inhibition.

### 8. N-acetylcysteine preserves the therapeutic efficacy of JIB-04 while mitigating hepatomegaly

We evaluated the effects of JIB-04 on peripheral organs, including the lung, brain, kidney, and liver. Among these tissues, only the liver exhibited reversible hepatomegaly accompanied by lipid accumulation (Fig. 8A, supplemental figure 1A and 1B). To further characterize this phenotype, we analyzed serum chemistry data from previous studies of tumor-bearing mice treated with JIB-04 or vehicle. JIB-04 treatment was associated with a mild increase in ALT and trends toward elevated cholesterol and triglyceride levels, consistent with hepatic steatosis and hepatocyte stress or injury (Supplemental figure. 1C).

**Figure 8.**
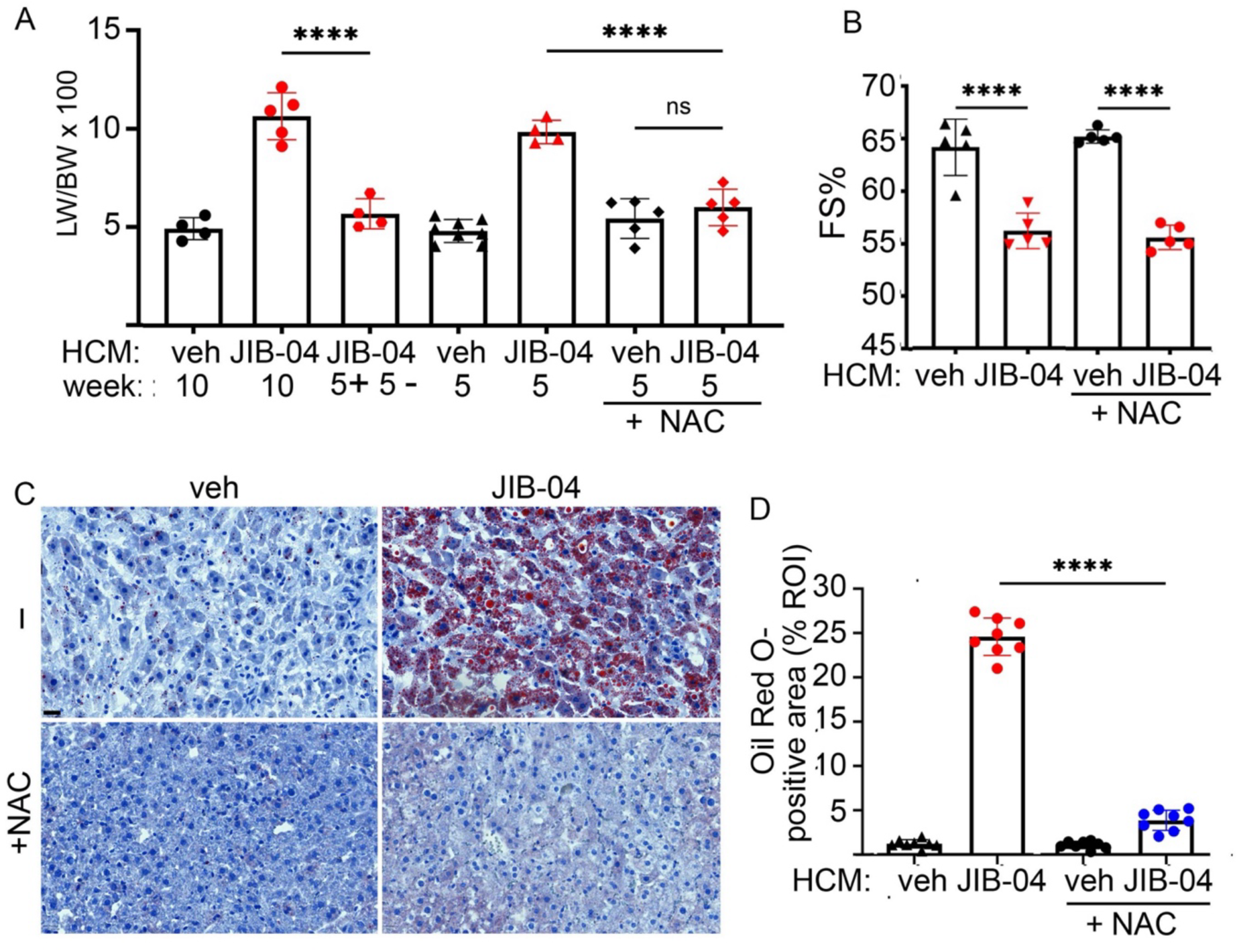
N-acetylcysteine (NAC) mitigates JIB-04-induced hepatomegaly and hepatic lipid accumulation while preserving anti-HCM efficacy. (A) Liver weight-to-body weight ratio (LW/BW) in HCM mice treated with vehicle, JIB-04 for 10 weeks, JIB-04 for 5 weeks followed by a 5-week drug withdrawal period (5+5−), JIB-04 for 5 weeks, vehicle plus NAC, or JIB-04 plus NAC. NAC significantly reduced JIB-04-induced hepatomegaly. (B) Fractional shortening (FS%) in HCM mice treated with vehicle or JIB-04 in the presence of NAC for 5 weeks. NAC did not impair the anti-HCM efficacy of JIB-04. (C) Representative Oil Red O staining of liver sections from mice treated with vehicle or JIB-04 in the absence or presence of NAC for 5 weeks. (D) Quantification of Oil Red O-positive area. JIB-04 induced marked hepatic lipid accumulation, which was substantially attenuated by NAC co-administration. Data are presented as mean ± SD. ****, p < 0.001.

To gain insight into potential mechanisms underlying this liver phenotype, we re-analyzed liver microarry data generated from a separate prior study of tumor-bearing mice treated with JIB-04 or vehicle. Differentially expressed genes were enriched in pathways related to xenobiotic metabolism, lipid metabolism, and inflammatory signaling. Consistent with activation of an adaptive xenobiotic response, genes encoding Phase I, II, and III detoxification enzymes—including cytochrome P450 (CYP) enzymes, glutathione S-transferases (GSTs), UDP-glucuronosyltransferases (UGTs), and ATP-binding cassette (ABC) transporters—were markedly upregulated (Supplemental figure. 2). Despite induction of these detoxification pathways, genes involved in hepatic lipid handling and lipid export remained dysregulated, suggesting incomplete adaptation to JIB-04-induced metabolic stress. Because oxidative stress is a common consequence of xenobiotic metabolism and can promote hepatic lipid accumulation, we hypothesized that it contributes to JIB-04-induced hepatomegaly.

To test this hypothesis, we evaluated co-administration of the antioxidant N-acetylcysteine (NAC) in the Myh6^R403Q/+^ HCM model. NAC co-treatment significantly attenuated JIB-04-induced hepatomegaly (Fig. 8A) while preserving the therapeutic efficacy of JIB-04 against HCM (Fig. 8B). In addition, NAC reduced hepatic lipid accumulation compared with JIB-04 treatment alone, as evidence by decreased Oil Red O staining (Fig. 8C and 8D). Together, these findings identify oxidative stress as an important contributor to JIB-04-induced hepatomegaly and suggest that antioxidant co-therapy may improve the tolerability of KDM inhibition while maintaining its therapeutic benefit.

## Discussion

Increasing evidence indicates that sarcomere mutations in HCM induce energetic and mitochondrial stress, triggering sterile inflammation and progressive myocardial remodeling ^40–42^. Consistent with this concept, we observed progressive macrophage activation, inflammatory signaling, and fibrosis at both the transcriptional and cellular levels during HCM progression in the CsA-accelerated Myh6^R403Q/+^ model. In the present study, pharmacological inhibition of histone lysine demethylases (KDMs) with JIB-04 prevented disease onset, reversed established pathological remodeling, and improved cardiac function in both accelerated and naturally progressing Myh6^R403Q/+^ mouse models. Mechanistically, JIB-04 restored disease-associated transcriptional and chromatin remodeling, identified PHF2 as a candidate component of a KDM regulatory network involved in HCM pathogenesis, and reproduced these molecular and functional effects in human MYH7 R403Q iPSC-derived cardiomyocytes, supporting the translational relevance of KDM inhibition. Collectively, these findings support a model in which sarcomere mutation-induced hypercontractility promotes mitochondrial dysfunction and DAMP release, leading to macrophage activation, inflammation, fibrosis, and progressive cardiac remodeling, while KDM inhibition interrupts this pathogenic cascade by reversing disease-associated epigenetic remodeling across multiple cardiac cell types and stages of disease progression.

An important finding of this study is that chromatin remodeling precedes overt pathological remodeling in HCM. ATAC-seq revealed widespread alterations in chromatin accessibility in non-diseased Myh6^R403Q/+^ hearts before the development of hypertrophy, fibrosis, or cardiac dysfunction, indicating that the pathogenic R403Q mutation initiates epigenetic reprogramming long before clinical disease becomes apparent. These early changes were characterized by reduced promoter accessibility together with redistribution of accessible chromatin toward distal regulatory regions, suggesting altered transcriptional control at the earliest stages of disease. As HCM progressed, these chromatin alterations were maintained and accompanied by activation of inflammatory, fibrotic, and stress-response transcriptional programs. Importantly, JIB-04 partially restored chromatin accessibility toward the WT pattern, supporting the concept that disease-associated epigenetic remodeling remains reversible even after pathological remodeling has been established. The substantial overlap between pathways identified by ATAC-seq and bulk RNA-seq further suggests that altered chromatin accessibility contributes directly to the transcriptional reprogramming that drives HCM progression. Together, these findings identify epigenetic remodeling as an early pathogenic event in HCM and provide a mechanistic basis for therapeutic intervention before irreversible structural remodeling occurs.

Among the candidate JIB-04 targets identified, PHF2 emerged as a particularly compelling mediator. PHF2 has previously been shown to undergo PKA-dependent activation and to demethylate H3K9me2, thereby promoting transcription of stress-responsive genes. Identification of PHF2 together with PRKACA and AKAP9 in biotin-JIB-04 pull-down experiments suggests that JIB-04 may regulate a PKA-PHF2 signaling axis that becomes activated during HCM. Although additional studies are needed to define the direct molecular targets of JIB-04, PHF2 knockdown in cardiomyocytes, macrophages, and fibroblasts phenocopied several effects of JIB-04, supporting an important contribution of PHF2 to pathological remodeling.

Our findings further indicate that HCM is associated with coordinated activation of multiple histone lysine demethylases and that the therapeutic effects of JIB-04 likely extend beyond inhibition of PHF2 alone. Although PHF2 was the only KDM identified by biotin-JIB-04 affinity purification, transcriptomic analyses of murine HCM together with published human single-nucleus RNA-sequencing datasets demonstrated coordinated dysregulation of multiple KDMs, including PHF2, KDM4A, KDM5A, and KDM6B, in genetically heterogeneous HCM hearts. The absence of additional KDMs in the pull-down experiments likely reflects technical limitations of affinity capture from nuclear lysates rather than exclusion of other functional targets. Importantly, PHF2 was identified in both mouse HCM hearts and human HCM septal tissue representing mixed genetic etiologies, and JIB-04 restored conserved transcriptional programs in both murine HCM and human MYH7 R403Q iPSC-derived cardiomyocytes. Together, these observations suggest that dysregulation of KDM-dependent epigenetic programs represents a common downstream consequence of diverse HCM-causing mutations rather than a mutation-specific event. Although additional studies in other genetic HCM models are warranted, our findings support KDM inhibition as a potentially mutation-agnostic therapeutic strategy targeting shared mechanisms of pathological remodeling.

Although JIB-04 produced reversible hepatomegaly accompanied by hepatic lipid accumulation, no overt toxicity was observed in other major organs. Analysis of liver transcriptomic data from an independent study of JIB-04-treated tumor-bearing mice revealed robust activation of xenobiotic detoxification pathways together with persistent dysregulation of genes involved in lipid metabolism and lipid handling, suggesting an adaptive but incomplete response to chronic JIB-04 exposure. Importantly, co-administration of N-acetylcysteine substantially reduced hepatomegaly while preserving the anti-HCM efficacy of JIB-04, supporting oxidative stress as an important contributor to the hepatic toxicity associated with KDM inhibition.

The molecular basis of this toxicity remains to be defined. One potential mechanism is that inhibition of PHF2 or other JIB-04-sensitive KDMs impairs adaptive antioxidant responses that normally protect hepatocytes during xenobiotic stress. Previous studies have shown that PHF2 can promote NRF2-dependent cytoprotective programs under conditions of metabolic stress,^43^ raising the possibility that disruption of this pathway contributes to hepatic lipid accumulation during chronic JIB-04 treatment. Although this hypothesis requires direct experimental validation, our findings identify oxidative stress as a modifiable component of the principal toxicity observed in this study and suggest that antioxidant co-therapy may improve the tolerability of KDM inhibition while maintaining therapeutic efficacy.

In conclusion, this study identifies KDM inhibition as a promising disease-modifying therapeutic strategy for HCM. By simultaneously targeting cardiomyocyte dysfunction, inflammation, fibrosis, and epigenetic remodeling, JIB-04 produced sustained therapeutic benefits across murine and human HCM models. These findings provide a mechanistic framework for the development of next-generation KDM inhibitors with improved selectivity and safety for the treatment of HCM.

## Acknowledgement

This work was supported by the National Institutes of Health/National Heart, Lung, and Blood Institute grants R01HL159599 (J.L.), R01HL145298 and P01HL160488 (D.J.C.), and R01HL157050 (Z.P.L.). E.D.M. was supported by the Robert A. Welch Foundation (I-1878). E.N.O. was supported by the National Institutes of Health (R01HL157281 and P01HL160488), the Robert A. Welch Foundation (I-0025), the Leducq Foundation Transatlantic Network of Excellence (20CVD04), and the British Heart Foundation Big Beat Challenge award to CureHeart (BBC/F/21/220106).

## Supplemental materials

**Supplemental Figure 1.**
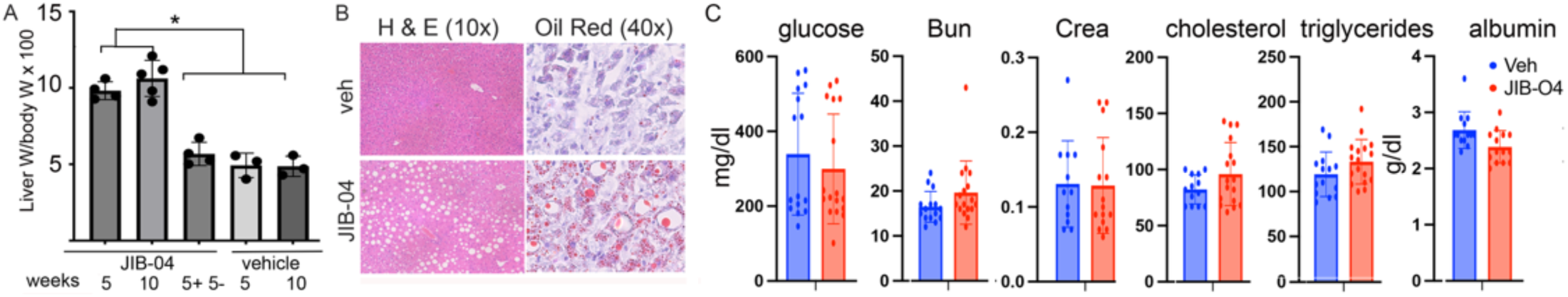
JIB-04 induces reversible hepatomegaly and hepatic steatosis. (A) Liver weight normalized to body weight in *Myh6^R403Q/+^* mice treated with vehicle or JIB-04 for 5 or 10 weeks or with JIB-04 for 5 weeks followed by a 5-week drug-free recovery period (5+5–), as outlined in Figure 3A and 4A. (B) Representative H&E (left) and Oil Red O (right) staining of livers from HCM mice treated with vehicle or JIB-04 for 10 weeks. (C) Serum metabolic analyses revealed no significant changes in glucose, BUN, or creatinine levels following JIB-04 treatment. Albumin levels were modestly reduced, whereas cholesterol and triglycerides exhibited trends toward elevation.

**Supplemental figure 2.**
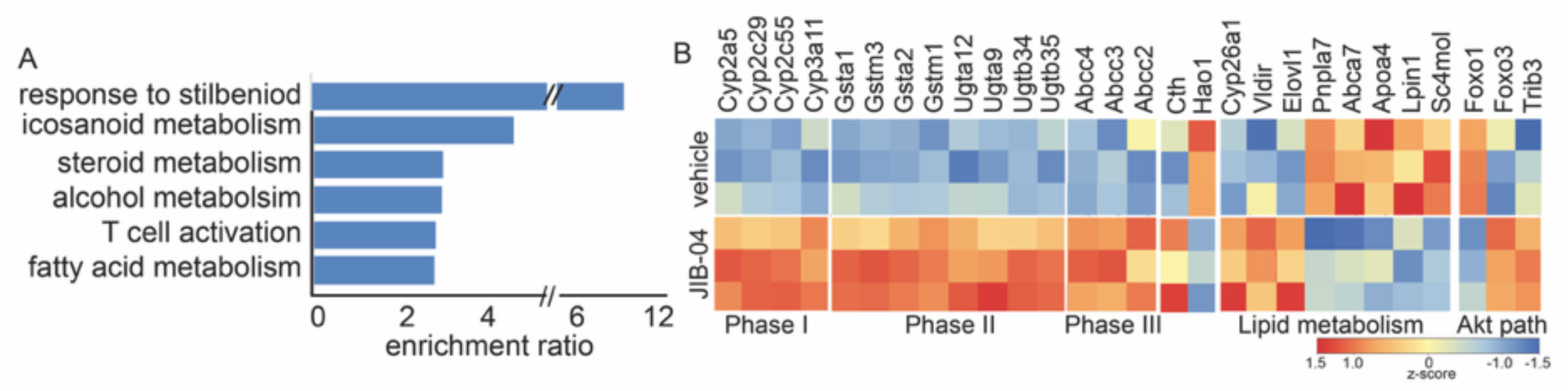
Transcriptomic changes associated with JIB-04-induced liver toxicity. (A) GO enrichment analysis of DEGs in livers from vehicle- and JIB-04-treated mice. (B) Heat map of representative DEGs altered by JIB-04 treatment, grouped according to functional categories including Phase I, II, and III detoxification pathways, lipid metabolism, and stress-response signaling.

